# Estimating Northern Fur Seal Pup Production: A Case Study of Batch Mark Abundance Estimation

**DOI:** 10.1101/772350

**Authors:** Devin S. Johnson, Rod G. Towell, Jason D. Baker

**Author notes:** Correspondence: Devin Johnson, Pacific Islands Fisheries Science Center, National Marine Fisheries Service, Honolulu, Hawaii 96818.

## Abstract

We describe a hierarchical N-mixture model for estimating northern fur seal pup production from batch mark-resight data. Our goal was to improve upon a traditional design-based estimation method used for over 50 years. To this end, we propose a hierarchical N-mixture model to account for differences in animal availability for resighting and observer detection probabilities. A Bayesian approach is used for inference with three separate methods proposed for necessary computations. First a straightforward posterior sample is drawn using MCMC. This was considered the gold standard for this analysis. However, we also consider an approximate model-based on Gaussian approximation of the Poisson and binomial distributions used in the exact hierarchical model. By using the Gaussian approximations, analytic integration can be used to marginalize over latent components. Inference can then be made by maximizing the posterior to find the mode. Following this we investigate both delta-method and parametric bootstrap approaches for calculating abundance and the associated standard errors with batch sampling data collected on northern fur seals in 2016. Each of the three newly proposed methods produced virtually identical estimates and standard errors. In addition, the parametric bootstrap approximation to a posterior sample was also virtually the same as the MCMC derived sample. There were a handful of small differences in the point estimates of abundance but these were not large enough to be especially notable. There were some noticeable differences in uncertainty estimates. The production estimate coefficients of variation ranged from 2.5–9.5% for the model-based estimates, but 0.0–12.5% for the design-based estimates. The design-based estimates of uncertainty were more volatile using the design-based method with some standard error estimates unreasonably small. Through a case study we have provided support for using Gaussian approximation in *N*-mixture type models when abundance is relatively large. These situations are typically challenging for fitting models using any sort of marginalization because there is no analytic solution for the exact models. However, when abundances are large using a Gaussian approximated model allows for analytic integration over latent abundance states which can greatly facilitate methodology such as maximum likelihood or posterior inference.

## 1 Introduction

Estimation of pup production has been the cornerstone for population and management decisions for the Pribilof Islands herd of northern fur seals (*Callorhinus ursinus*; NFS) for over 50 years. The design-based method of estimating production has remained largely unchanged since the late 1960s. Here we consider a modern hierarchical modeling approach for abundance estimation that allows for day-to-day differences in the proportion of pups that are available to be counted, as well as different rates of detection among observers. Although we do not make use of it here, the model-based version allows for use of covariates might be helpful in the future for modeling differences in parameters like detection probabilities or allow for addition of prior information on the proportion of animals that are marked. Although this method was designed and applied to NFS pup abundance, it is a general mark-resight modelling approach that could be used for many other species.

Since the mid-1960s, estimation of NFS pup production in the Pribilof Islands, Alaska has been accomplished using a design-based mark-resight method where pups are batch marked and resighted on at least 2 later dates. Chapman and Johnson (1968), York and Towell (1996), and Towell et al. (2016) provide detailed descriptions of the marking, resight, and current production estimation methods. In the first half of August, pups are batch marked by shearing the guard hairs on top of the head to make the light underfur conspicuous to observers. The light-colored underfur represents the mark. The goal of shearing is to mark approximately 10% of the pups on each rookery. The 10% is based on the previous estimate of pup production for the sample rookeries (typically 2 years prior). Shear marks are allocated proportionally on each rookery by section according to the fraction of the total number of breeding males counted in each section. Thus, the marks are spatially allocated to roughly match spatial abundance of pups. Marked and unmarked pups on each rookery are counted by two observers scanning (with the aid of binoculars when necessary) on at least two occasions.

Here we briefly describe the current design-based method of estimation. It is based on the cluster sampling method outlined by Chapman and Johnson (1968); however, the cluster groups are kept fixed at 25 pups, therefore the estimator formulas can be reduced to those below. In the current method, each resight day is considered an independent replicate. For each replicate *j* = 1, 2, a mark-frequency estimate is calculated, 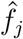 (total marks counted / total pups observed), the frequency of marked pups to total pups counted on the *j*th occasion. From this frequency estimate the occasion-specific estimate of the number of pups alive at the time of shearing is estimated as,

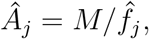

where *M* is the total number of pups marked. Note that this occasion specific estimate is identical to the classic Lincoln-Petersen estimator (Lincoln, 1930). However, the sampling procedure is different with pups often being counted twice by the two different observers, thus, a different standard error (SE) method is used. The full estimate is calculated by *Â* = *mean*(*Â_j_*). The standard error of this estimate can be estimated using the fact that 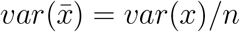. Thus, for 2 resight occasions, this becomes

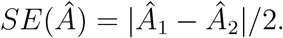

Because the *SE*(*Â*) is based on only two replicated samples the estimate can become unstable, often producing estimates that are unreasonably small. See the Results section for example.

In addition to mark-resight methods that are applied to live pups, dead pups are counted on a sample of rookeries. Each sampled rookery is completely examined and the number of dead pups are recorded. For these sampled rookeries, the number of dead pups is added to obtain the pup production estimate 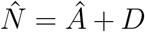. For those rookeries 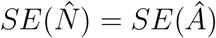. However, for rookeries without dead pup samples, 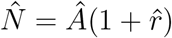 where the ratio of dead pups to live pups, 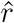, is estimated using a ratio estimation method (Cochran, 1977) based on the rookeries where dead pups are counted.

In this paper we propose an updated, model-based methodology for analyzing the same batch mark data that are currently collected. The model-based approach is similar in spirit to the open N-mixture batch sampling models of Cowen et al. (2017) but we consider the populations to be closed over the study, however, different proportions of the population are unavailable for sampling in each resight occasion. Thus, abundance does not follow the open population models of Dail and Madsen (2011), but rather in each resight occasion animals have a probability of not being detectable at all. This may occur if they are in the water or not visible from sampling positions. In addition to availability, the model-based approach also allows explicit modeling of different detection rates between observers. Finally, with the model-based approach we can directly model neonate mortality simultaneously with the mark-resight data without the need to use the additional bull count data to separately estimate dead pup numbers. Although we do not explore it here, by explicitly modeling neonate mortality, we also introduce the possibility of modeling this parameter with environmental covariates that may be useful in determining what conditions lead to increased (or decreased) neonate mortality.

We begin development in the following section describing the model pieces in a hierarchical version of an N-mixture model (Royle, 2004) that accounts for differential availability of animals during each resight occasion. Next, we propose two different methods of Bayesian estimation for abundance and model parameters. The first is a straightforward implementation in the Markov chain Monte Carlo (MCMC) software `JAGS` (Plummer, 2003).

In the second approach we consider an asymptotic approximation similar to Brintz et al. (2018) where the latent discrete components of the model are treated as continuous valued and modeled with a multivariate normal distribution (Normal). This is motivated by the fact that the unknown abundance is large and little is lost empirically but much is gained analytically by this approximation. The unobserved discrete components can be marginalized out of the model to improve numerical procedures of estimation. In addition, one can also use automatic differentiation-based software such as `stan` (Gelman et al., 2015) and Template Model Builder [`TMB`; Kristensen et al. (2016)] which cannot fit models with discrete latent components directly. Both of these approaches are used to fit data collected in 2016 by the National Marine Fisheries Service as part of the biannual monitoring survey of Pribilof Islands pup production.

## 2 Methods

Pup production, *N*, is defined as the combined total of the number of pups alive at the time of the marking study, *A*, as well as, the number of pups that have died before marking, *D*. The notation for the descriptions in this section are summarized in Table 1 for further reference. Notably, *i* indexes a rookery (or site), *j* indexes a resight occasion, and *k* indexes an observer. The number of pups batch marked at each rookery is *M_i_*, the unobserved total number of unmarked pups is *U_i_*, and the number of pups that died before the survey is *D_i_*, thus, pup production is *N_i_* = *M_i_* + *U_i_* + *D_i_*. For all rookeries *M_i_* is known and for *some* rookeries *D_i_* is observed via dead pup counts that take place after the resighting occasions. While this likely does not represent the full number of dead pups it helps to account for neonatal death in the final estimate (it is usually on the order of 2–5% of *N*). The number of marked and unmarked pups counted by the *k*th observer on the *j*th occasion is *m_ijk_* and *u_ijk_*, respectively.

**Table 1:** Glossary of terms for the mark-resighting of northern fur seals.

| Notation | Description |
| --- | --- |
| $i$ | Rookery (site) index ($i = 1, \dots, I$) |
| $j$ | Resight occasion index ($j = 1, \dots, J$) |
| $k$ | Observer within occasion index ($k = 1, \dots, K$) |
| $N_i$ | Total pup production ($N_i = M_i + U_i + D_i$) |
| $D_i$ | Number of pups that died before marking |
| $M_i$ | Number of pups batched marked |
| $U_i$ | Number of pups that are unmarked |
| $A_i$ | Number of pups alive at marking ($A_i = M_i + U_i$) |
| $M_{ij}$ | The number of marked pups available for resight on $j$th occasion |
| $U_{ij}$ | The number of unmarked pups available for resight on $j$th occasion |
| $m_{ijk}$ | Number of marked pups counted on resight effort $j$ by the $k$th observer |
| $u_{ijk}$ | Number of unmarked pups counted on resight effort $j$ by the $k$th observer |
| $\xi_i$ | Probability that a pup dies before batch marking |
| $\tau_i$ | Probability that a pup is marked given it is alive when marking occurs |
| $\alpha_{ij}$ | Probability that a pup can be observed at a resight occasion by an observer (*e.g.*, not in the water or hidden from view) |
| $\delta_{ijk}$ | Probability that a visible pup is counted at a rookery by an observer |

### 2.1 A hierarchical N-mixture approach

The model-based approach we develop here seeks to improve on the design-based method of the last section by accounting for variability in the detection process, as well as differing availability of animals to be observed between resighting occasions. This latent availability induces positive correlation between observers on any single occasion.

The top level of the hierarchical model begins with the usual N-mixture model choice of

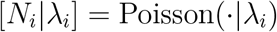

for the pup production model, where *λ_i_* is a rate hyperparameter. In all the description herein, we use the notation “[*x*]” and [*x*|*y*]” to represent the probability density function (PDF) of *x* and the conditional PDF of *x* given *y*. We do not necessarily have to use this model, but it can make it sufficiently uninformative (see following section).

The second level of the model partitions the *N_i_* pups born into three categories: (1) those that died before the survey, *D_i_*, (2) those that were marked at the start of the survey, *M_i_*, and finally, all those pups that were not marked, *U_i_*. Assuming that each pup is independently placed in 1 of the 3 categories, we can integrate over the unobserved *N_i_* to obtain the model

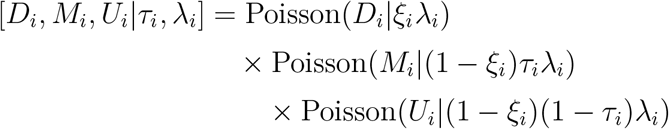

where *ξ* is the neonate mortality probability, and *τ_i_* is the probability that a pup is marked given it was alive at the start of the survey. Notice that we actually do not need to estimate *N_i_* as a parameter, because *M_i_* is known; we only need estimates of *U_i_* and any unobserved *Di* to estimate pup production.

Now that we have defined a model for (*M_i_, U_i_, D_i_*) we can examine the next level of the model which is the availability process for how many pups are available to be observed on subsequent resights. On occasion *j* the number of pups from each subpopulation that are observable (i.e., in the open and not in the water) is modeled by

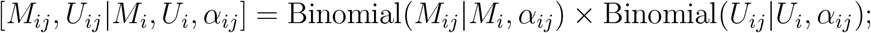

that is, each individual pup is randomly available for observation on each occasion. Given the number of pups available from each subpopulation, each observer randomly detects each pup in their counts. Thus for observer *k* on occasion *j* the data models for the observed counts are

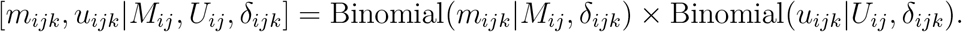

In the collection of the NFS pup production data here, great care is taken to avoid double counting the same individuals. The resighters methodically proceed through the rookery counting small groups that are not spatially overlapping. However, if it is not possible to assume that individuals will only be observed, at most, once during an occasion, then another distribution can be used here. For example, we might assume that marked and unmarked animals are encountered with a rate proportional to availability, thus,

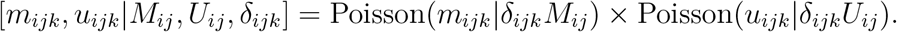

would be a good model for this situation. In the remainder of this paper, we will assume the binomial version for the observation model.

For each rookery, the complete likelihood for the hierarchical model is

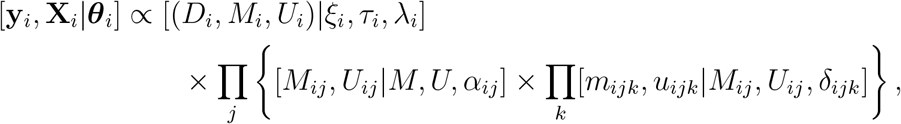

where ***θ**_i_* is used to denote all of the parameters such as *δ_ijk_* and *α_ij_*, **X***_i_* represents all the latent abundance values (*U_i_, M_ij_*, and *U_ij_*), and *y_i_* is the observable data (*M_i_, m_ijk_*, and *u_ijk_*). Note, that for some rookeries, *D_i_* will be included in **X**_*i*_ and for the others it will be in y*_i_*. Now, we can make inference to the latent quantities and parameters with the full complete-data likelihood,

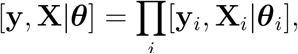

by taking the product over rookeries.

### 2.2 Bayesian inference

Traditional maximum likelihood estimation is difficult as the appropriate likelihood to maximize would be

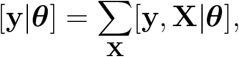

where the sum is over all possible values of the high-dimensional, unobserved **X**. Numerically calculating the integrated likelihood over every iteration of a maximization routine would be daunting. Cowen et al. (2017) illustrate an efficient numerical method for an open population model using a hidden Markov model formulation. However, the latent structure here is not time indexed in a way that makes HMM formulation straightforward.

The most straightforward way of estimating parameters and abundance from the model developed in the previous section is to sample from the posterior distribution using MCMC. The appropriate posterior distribution in this case

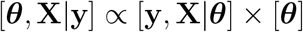

where [***θ***] is the prior distribution of the parameters.

For specification of the prior distribution we will begin with *λ_i_*. Link (2013) suggested that *N_i_*, should follow the prior distribution 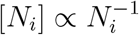, an improper prior. This can be approximated with the Poisson-gamma mixture

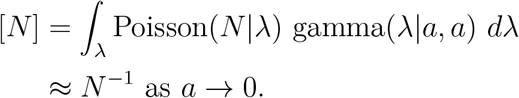

Thus, we suggest [*λ_i_*|*α*] = gamma(*λ_i_*|*a, a*) with *a* = 1.0e-6 (or something small). For the detection and availability parameters, we switch to the logit parameterization

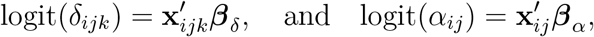

where **x** contains covariates such as weather conditions or observer related variables. They are not necessarily the same between the parameter types. A prior distribution is specified for each *β*; such as a multivariate normal distribution. The mortality and marking probabilities, *ξ_i_* and *τ_i_* could also be modeled this way, but for this investigation we leave their parameterization in its natural form and use beta or uniform prior distributions for those parameters.

### 2.3 An approximate hierarchical model

An alternative to working with the model as it stands in the “exact’’ form presented in the previous section is to analytically integrate over the latent components to produce the appropriate likelihood for the model and data. However, the latent components, *U_i_*, *U_ij_*, and *M_ij_* are all discrete with infinite upper bounds, thus the marginalization over these components involves many intractable infinite sums. Thus, straight forward maximum likelihood is difficult. The typical approach is to pick fixed upper bounds and execute the sums to those bounds (e.g., Dail and Madsen (2011), Schmidt et al. (2015), Cowen et al. (2017)). With large abundances this is still a monumental task. In the example of the next section the smallest rookery produces approximately 500 pups, so, the sums necessary become quite cumbersome.

In an attempt to overcome the computational issues of large abundances in open N-mixture models Brintz et al. (2018) propose a normal approximation that allows analytic integration to be performed, thus producing an asymptotically approximate integrated likelihood. Maximization of the likelihood can then be accomplished in a fraction of the time MCMC takes to run while still producing comparable results. In this section, we derive an asymptotically approximate likelihood for our batch marking capture-recapture model using the same principles.

The goal of this approach is to allow modeling the same data with an approximation of the general form

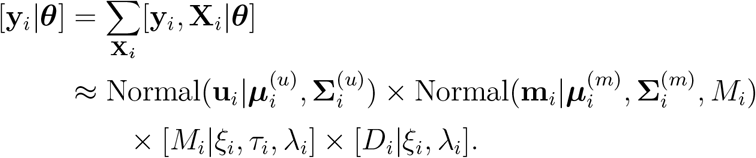

The approximation is accomplished by matching the mean vectors and covariance matrices to that of the marginal distribution of m*_i_* and u*_i_* after integrating over the latent components.

To obtain the multivariate normal means and covariance matrices in the approximation, we will use the same moment matching approach as Brintz et al. (2018). The following means and covariance are derived using the independence of *M_i_* and *U_i_* given the parameters and the law of total covariance. First, the mean vectors, are straightforward to produce;

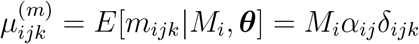

and

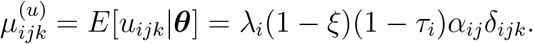

The covariance becomes a little more notationally challenging, but they can also be calculated using the standard unconditional variance formula,

*Cov*(*X, Y*) = *E*[*Cov*(*X,Y*|*Z*)] + *Cov*{*E*(*X*|*Z*), *E*(*Y*|*Z*)}. First we define some terms to make the notation easier to follow. We will use *ω* = (*i, j, k*) to index a specific site, occasion and observer. Next we define the three functions

- *σ_i_*(*ω, ω′*) = *α_ij_α_ij′_δ_ijk_δ_ij′k′_* for *i* = *i′* and 0 else-wise
- *σ_ij_*(*ω, ω′*) = *α_ij_*(1 – *α_ij_*)*δ_ijk_δ_ijk′_* for *i* = *i*′, *j* = *j*′ and 0 else-wise
- *σ_ijk_*(*ω, ω′*) = *α_ij_δ_ijk_*(1 – *δ_ijk_*) for *ω* = *ω*′ and 0 else-wise

Now, the entries of the covariance matrices can be specified as

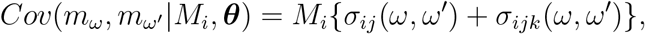

for *i* = *i′*, *j* = *j′*, and 0 else-wise. For the unmarked animals we are also marginalizing over *U_i_* as well, covariances are thus positive over different occasions; that is

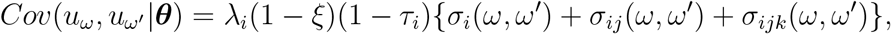

for *i* = *i′* and 0 else-wise.

Using this approximation, we can form the marginal posterior distribution,

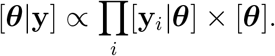

From this specification we can use MCMC to draw from the posterior distribution or use optimization methods to make simulation free inference. The major numerical argument for this approximation is that it marginalizes over all discrete valued components of the model. Therefore, the remaining parameters are all continuously valued, thus MCMC or optimization can be carried out with software that uses automatic differentiation such as `stan` or `TMB`. In addition, it is often note that marginalizing over some model components can improve posterior sampling or optimization (Van Dyk and Park, 2008).

Now that we have integrated **X**_*i*_ out of the models, it begs the question, how can we now estimate *N_i_* = *M_i_* + *U_i_* + *D_i_* when *U_i_* and some *D_i_* are no longer in the model? There are two approaches we can take, but they are both based on the posterior predictive distribution of the needed quantities. If *D_i_* is unobserved, then we can write down the predictive distribution

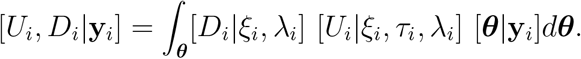

The obvious first approach for making abundance inference is to sample from these distributions by first drawing a sample from [***θ***|**y***_i_*] using, say MCMC, then for each draw, select a *D_i_* and *U_i_* from their respective Poisson distributions. However, we were seeking to produce a MCMC free method which might be computationally inexpensive. So, we can optimize [***θ***|**y**_*i*_] with respect to ***θ*** to obtain the posterior mode, then using the calculated posterior Hessian, **H**, further approximate

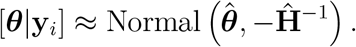

From this approximation one can readily draw thousands of samples in seconds. We will denote this approach as the parametric bootstrap method.

For an alternate fully simulation-free approach we can use the same technique as Johnson et al. (2010). That is, we can use the standard unconditional mean and variance formulas

- *E*(*X*) = *E_Y_* {*E_X_* (*X*|*Y*)}
- *V ar*(*X*) = *V ar_Y_* {*E_X_* (*X*|*Y*)} + *E_Y_* {*V ar_X_* (*X*|*V*)}

to approximate the mean (the optimal predictor) and variance of the predictive distribution for *U_i_* and *D_i_*. That is,

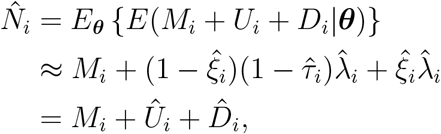

where the last term is replaced with *D_i_* if it is observed. The parameter estimates can be obtained by the posterior mode 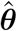 found by optimization as before. The prediction variance is calculated by

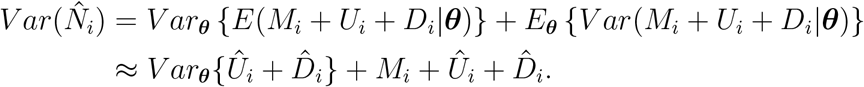

Because 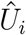 and 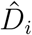 are just nonlinear functions of the parameters we can approximate the variance using the delta method (Dorfman, 1938) based on the asymptotic posterior variance, 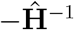 (Johnson et al., 2010).

There are benefits to both the bootstrap and delta method approaches. For the parametric bootstrap approach, the main benefit is the ease in which estimates of derived parameters can be calculated. As with the full MCMC one must only calculate the derived parameters for all the sample draws and one has a posterior sample of the derived parameter. In the next section, for example, we examine total pup production for each island. The delta method approach maintains the benefit that the calculations are quick and free of simulation issues such as choice of sample size, etc. In the following section, we examine how each of these approximate approaches compare to the “exact’’ MCMC approach, as well as, the traditional design-based estimates.

### 2.4 Pup production in 2016

Here we analyze the northern fur seal pup production data obtained in 2016 for monitoring pup production in the Pribilof Islands, Alaska (Figure 1). There are 19 separate rookeries on the two main islands in the archipelago, St. Paul Island in the north and St. George Island to the south. The rookeries on St. Paul Island produce approximately 80% of the pups and contains the majority of the rookeries. The 2016 data are fully described by Testa (2018). We built an R (R Core Team, 2019) package called `mbpp` that contains the model estimation procedures presented in the previous section and the data analyzed herein. This package is available at https://github.com/dsjohnson/mbpp and can be directly installed using the devtools package. The github page provides instructions for installation. We use the MCMC software `JAGS` (Plummer, 2003) through the `R` package `R2jags` (Su and Yajima, 2015) to sample from the exact hierarchical model and the R package `TMB` (Kristensen et al., 2016) to optimize the integrated posterior distribution for the approximate model bootstrap and delta approaches.

**Figure 1:**
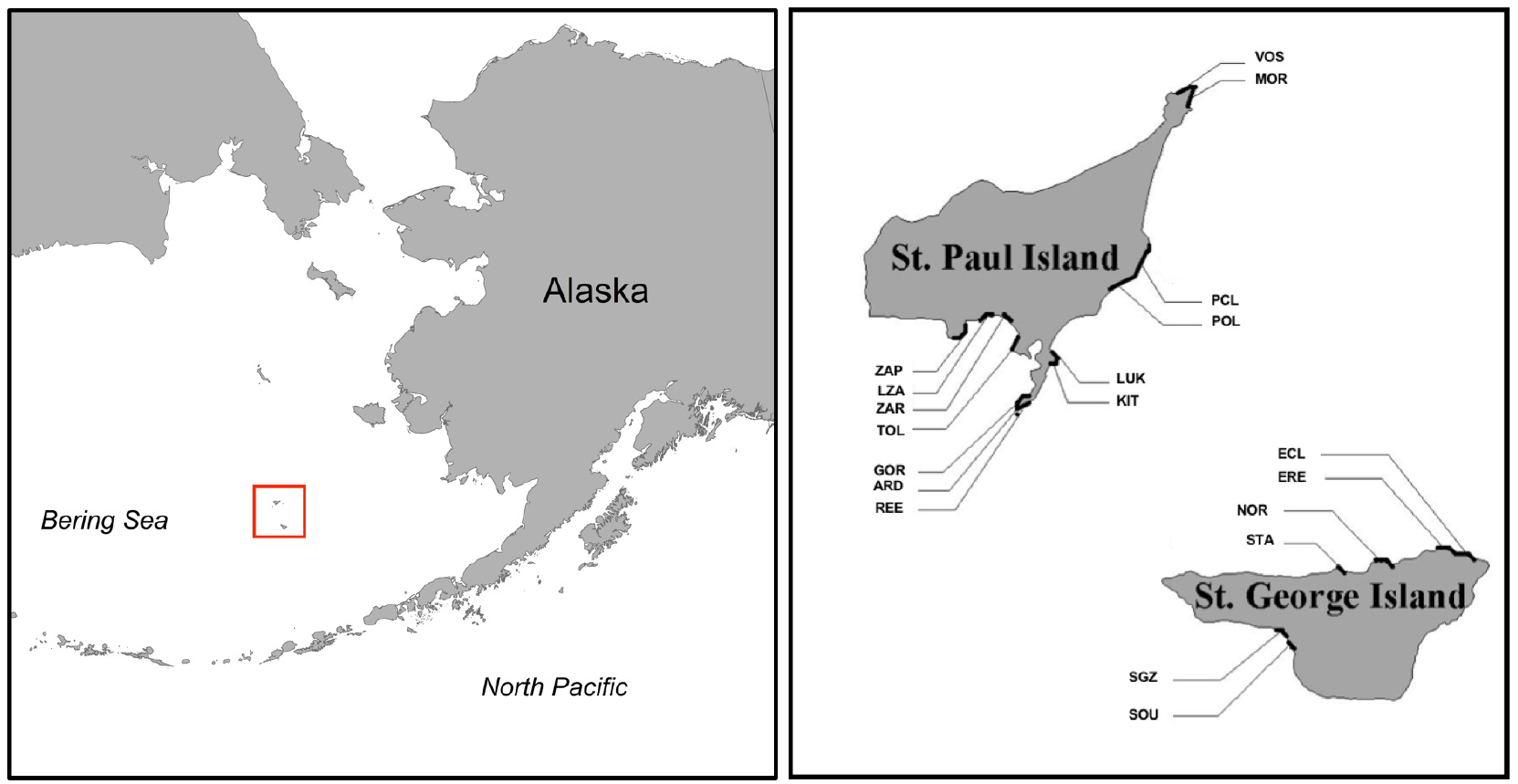
Map of northern fur seal rookeries in the Pribilof Islands, Alaska. The Pribilof Islands are an archipelago in the Bering Sea off the coast of Alaska. The main islands are St. Paul Island, to the north, and St. George Island, to the south. Note, the distance between the islands is not to scale in the right-hand figure. The rookeries are each labeled by their 3-letter NMFS rookery code.

For each of the rookeries (*I* = 19) in 2016, there were 2 observers (*J* = 2) who visited each rookery twice (*K* = 2) after shear-marking to count marked and unmarked pups. Each of the 3 model fitting methods presented in the previous sections was used to estimate pup production. Parameterizations and prior distributions are summarized in Table 2 for reference. For this analysis we used the previously suggested gamma distributions for each rookery to approximate a scale prior distribution for *N_i_*. Neonate mortality was modeled as constant within each island so the number of dead pups at unsampled rookeries could be predicted. The proportion of shear-marked pups (*τ_ij_*) was separately modeled for each rookery with a uniform prior. For the availability (*α_ij_*) and detection (*δ_ijk_*) we used a logistic regression parameterization to model rookery, occasion, and observer differences. The coefficients for each term were treated as normal random effects. So, for each term in the model, a variance component was estimated. This allows for regularization of the coefficients to be used for model selection (Hooten and Hobbs, 2015). An exponential prior distribution was used for each of the variance parameters to penalize complexity (Simpson et al., 2017).

**Table 2:**
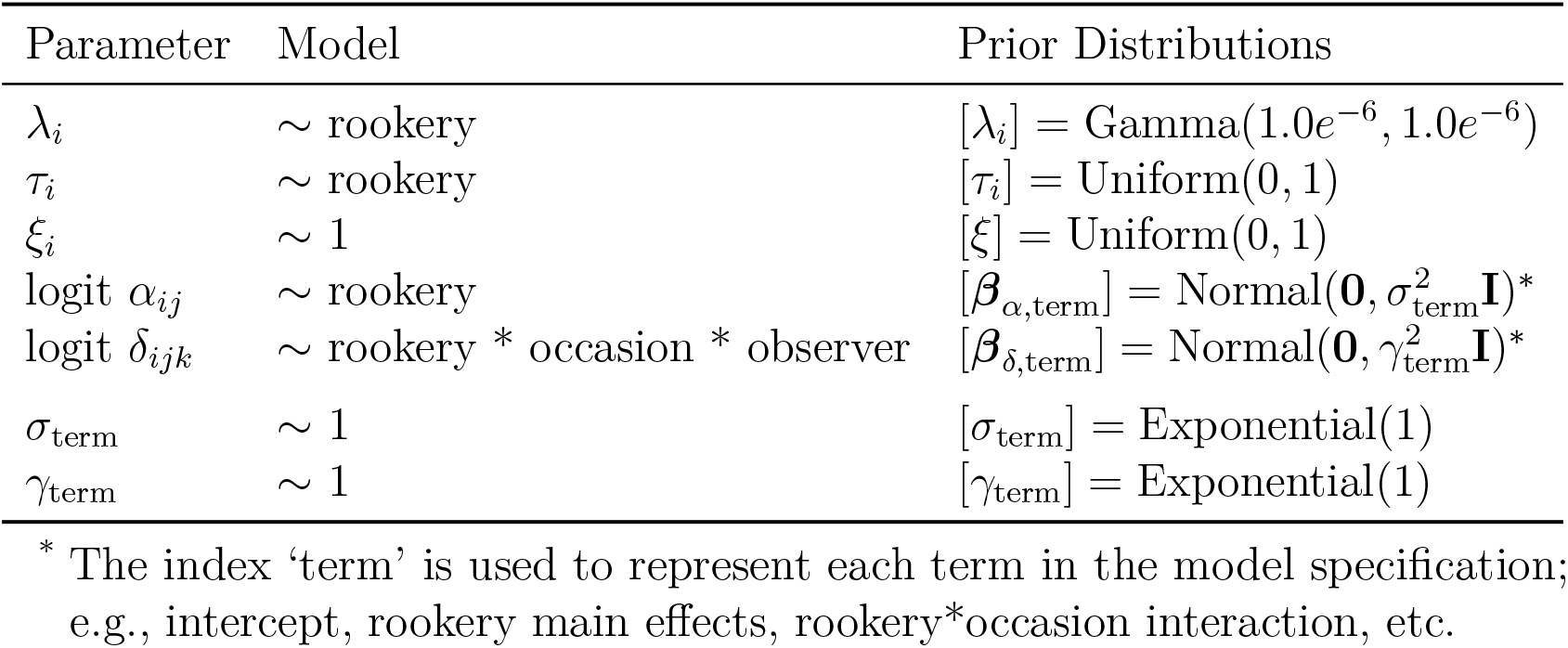
Parameter formulation and prior distribution specification. The ’Model’ column uses R notation to specify variation in the parameters over rookeries, occasions, and observers. Pup production was estimated separately for each island, therefore an intercept only model (~ 1) would mean that the parameter is constant within each island.

| Parameter | Model | Prior Distributions |
| --- | --- | --- |
| $\lambda_i$ | ∼ rookery | $[\lambda_i] = \text{Gamma}(1.0e^{-6}, 1.0e^{-6})$ |
| $\tau_i$ | ∼ rookery | $[\tau_i] = \text{Uniform}(0, 1)$ |
| $\xi_i$ | $\sim 1$ | $[\xi] = \text{Uniform}(0, 1)$ |
| logit $\alpha_{ij}$ | ∼ rookery | $[\beta_{\alpha, \text{term}}] = \text{Normal}(\mathbf{0}, \sigma_{\text{term}}^2 \mathbf{I})^*$ |
| logit $\delta_{ijk}$ | ∼ rookery * occasion * observer | $[\beta_{\delta, \text{term}}] = \text{Normal}(\mathbf{0}, \gamma_{\text{term}}^2 \mathbf{I})^*$ |
| $\sigma_{\text{term}}$ | $\sim 1$ | $[\sigma_{\text{term}}] = \text{Exponential}(1)$ |
| $\gamma_{\text{term}}$ | $\sim 1$ | $[\gamma_{\text{term}}] = \text{Exponential}(1)$ |

\* The index 'term' is used to represent each term in the model specification; e.g., intercept, rookery main effects, rookery\*occasion interaction, etc.

For each island, the MCMC algorithm was run for 100,000 iterations following a burnin of 10,000 iterations. Every 100th iteration was retained to obtain a final sample of 1,000 posterior draws. In order to fit the approximate model, we used Laplace approximation functionality in `TMB` to numerically integrate over the random coefficients and maximize the marginal likelihood. Then we used the delta-method and parametric bootstrap approaches for inference.

## 3 Results

The overall conclusion of the separate analysis of the shear sampling data using the 3 proposed methods in this paper revealed very little difference in the results between the model-based estimates. Certainly, there were no qualitative or management-level differences between the three estimation procedures considered here (Figure 2). In addition, there was no appreciable difference in execution time. All three methods finished in a little less than 2 seconds. However, this data set is relatively small, so there may be a larger differences with larger data sets.

**Figure 2:**
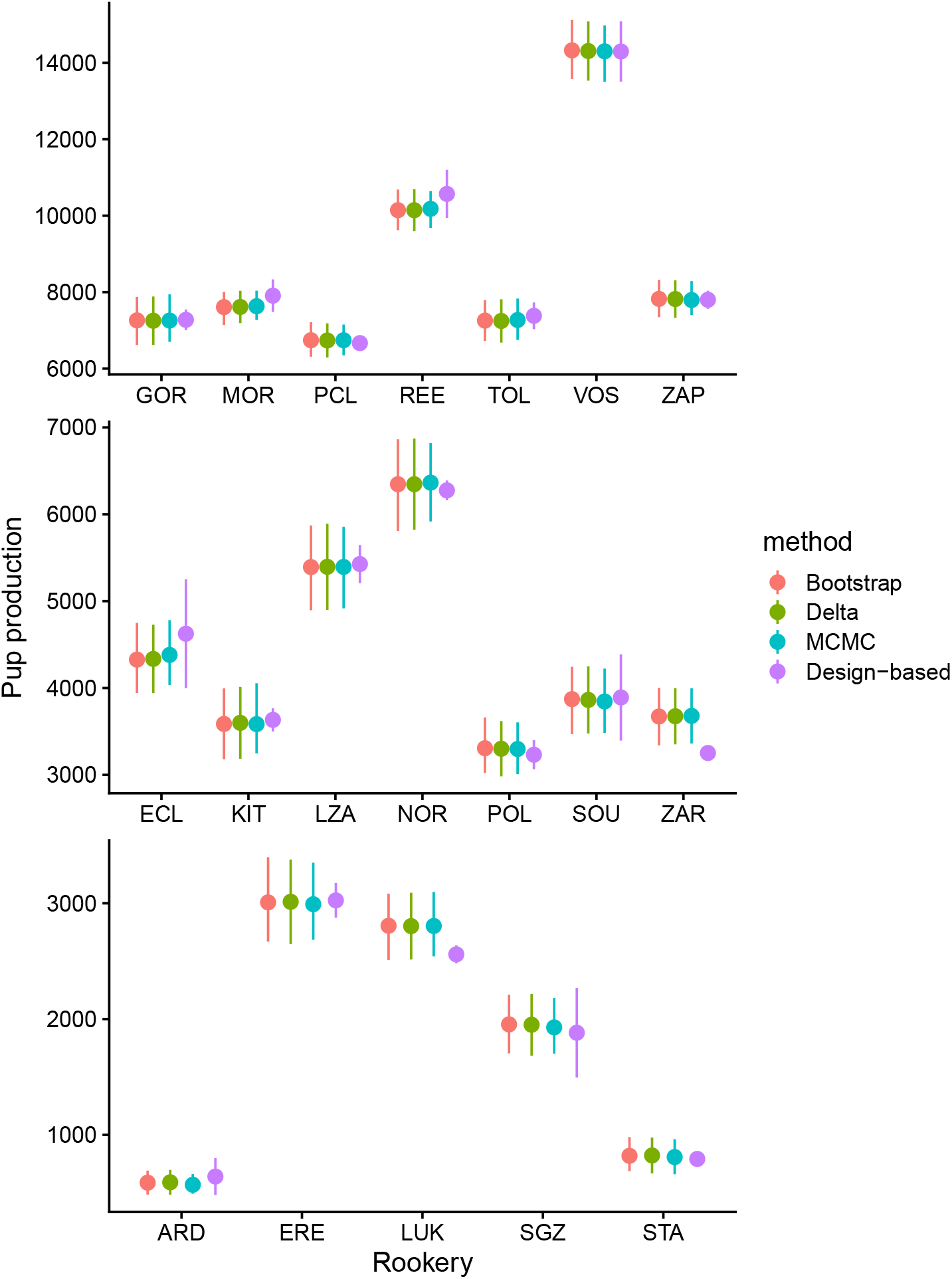
Estimated northern fur seal pup production. Each color represents a different model fitting method. Blue is a full MCMC sampling of the hierarchical model, red in the asymptotic normal approximation fitted with Template Model Builder using Laplace approximation, green in the asymptotic normal approximation with estimation based on a Gaussian parametric bootstrap, and purple is the traditional design-based estimator. The figure is divided into general rookery size classes for presentation.

There were some differences between the design-based and model-based point estimates, for example ZAR and LUK (Figure 2). However, these differences are not significant in terms of possible management decisions. There were some noticeable differences in uncertainty estimates. The production estimate coefficients of variation ranged from 2.5–9.5% for the model-based estimates, but 0.0–12.5% for the design-based estimates. The design-based estimates of uncertainty were more volatile using the design-based method with some standard error estimates unreasonably small. For example, production at PCL was estimated to be on the order of 7,000 pups, but the design-based SE was estimated to be 0.5! The model-based estimators produced an SE around 215 for PCL, much more reasonable.

To examine whether we might see differences between the bootstrap method versus the MCMC sample for estimating derived parameters we also estimated total pup production for each island. These island total estimates are the pup production quantities typically reported by NMFS for management decisions. We only compared the bootstrap and MCMC methods because they are both Monte Carlo-based methods which allow straightforward estimates of derived parameters. Both methods produced very similar point and interval estimates (Table 3). In addition, the distribution of the island total samples were also very similar (Figure 3). So, in this instance, the bootstrap approach seems to produce a nearly identical posterior sample compared to MCMC sampling of the exact hierarchical model. The model-based methods produced very similar point estimates when compared to the design-based method. The SE estimate was slightly higher for the model-based estimates due to the fact that the design-based SE was unreasonably small for some rookeries. However, the overall difference was not great (Table 3).

**Table 3:** Estimates of total pup production for each island using MCMC sampling of the exact hierarchical model, parametric bootstrap samples from the approximate model, and the traditional design-based estimator.

| Method | N | SE | CI |
| --- | --- | --- | --- |
| **St George** |  |  |  |
| MCMC | 20,356 | 485 | [19,449 – 21,359] |
| Bootstrap | 20,354 | 494 | [19,398 – 21,309] |
| Design-based<sup>1</sup> | 20,490 | 454 | [19,395 – 21,646] |
| **St Paul** |  |  |  |
| MCMC | 80,533 | 852 | [79,044 – 82,263] |
| Bootstrap | 80,526 | 906 | [78,816 – 82,269] |
| Design-based<sup>1</sup> | 80,641 | 717 | [79,135 – 82,176] |

<sup>1</sup> Design-based estimates taken from Testa (2018) “Population assessment of northern fur seals on the Pribilof Islands, 2015–2016.”

**Figure 3:**
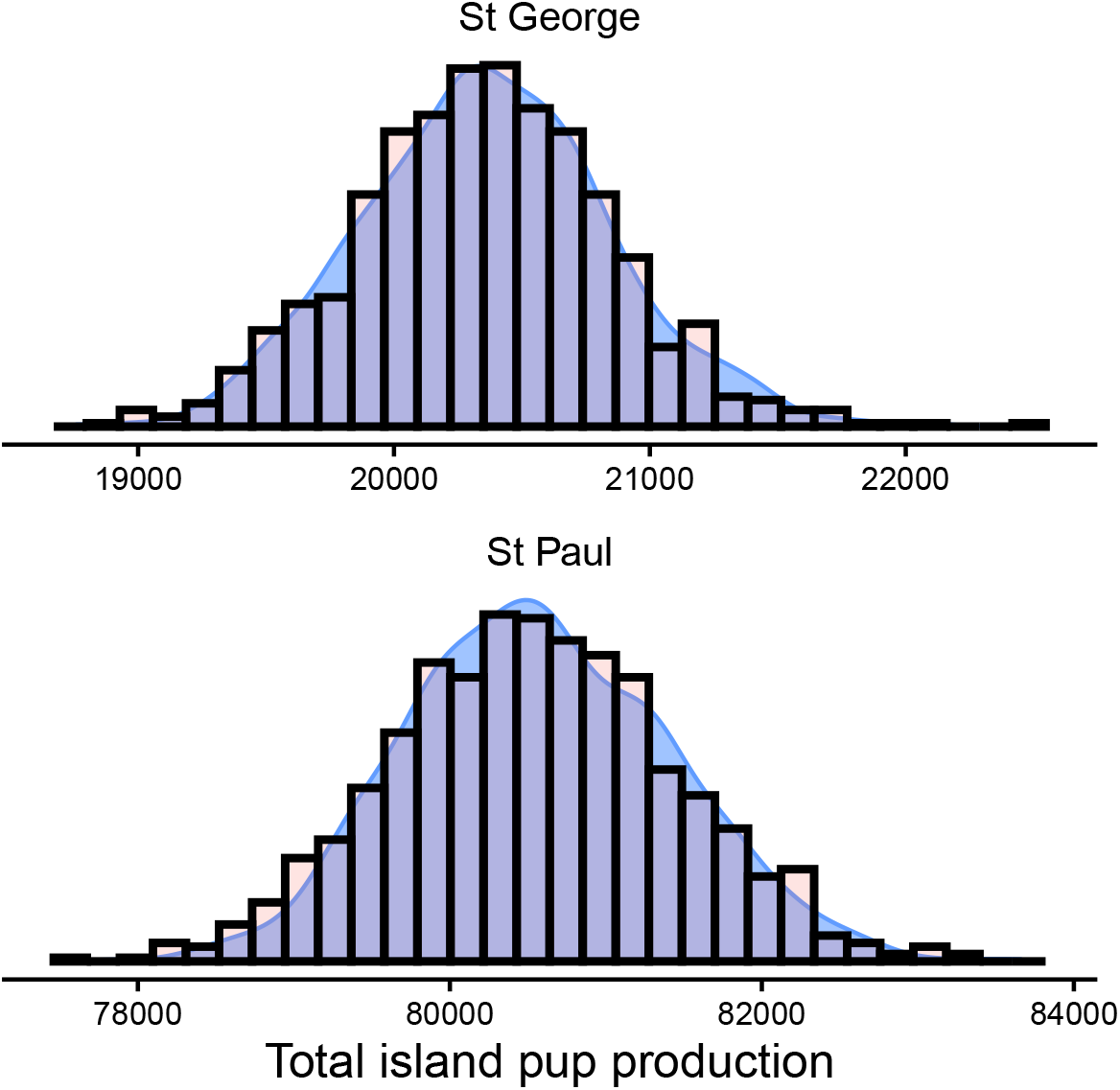
Estimated island-wide total northern fur seal pup production. The histogram represents the parametric bootstrap sample using the asymptotic approximation model. The density plot comes from the MCMC sample drawn from the exact hierarchical model using `JAGS`.

## 4 Discussion

Here we considered a model-based approach to estimating pup production for the Pribilof Islands population of northern fur seals. In the end, the model-based version produced similar point estimates, but more stable standard error estimates. With the model-based approach there is also a floor on the uncertainty estimation of *U_i_*, even if *τ_i_* and *ξ* are perfectly known, there is still binomial variation present in the model for *U_i_*. Therefore, unlike the design-based method, 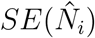 will never be zero because it is the prediction variance for an unobserved variable. Thus, producing realistic uncertainty assessment for each rookery.

Although there were not great computational savings, it was quite encouraging that the continuous-value approximate hierarchical model produced very similar results to the MCMC sampling of the exact model. Brintz et al. (2018) also noticed little difference in approximating the count data of open N-mixture models with a continuous valued multivariate normal distribution. This might form a blueprint for use of continuous approximation for open-population batch mark models such as those considered by Cowen et al. (2017) or the known-fate assisted N-mixture models proposed by Schmidt et al. (2015). In general, using mean and covariance matched Gaussian distributions may prove to be a useful approach for modeling latent discrete-valued variables in automatic differentiation based software such as `TMB` and `stan`. These models are ubiquitous in abundance estimation.

In addition to improvements over the design-based methods, the model-based method adds room for improvement for other ecologically interesting questions. As mentioned previously, we might choose to model *α_ij_* in some interesting way. For example, a temporal trend to reflect the increasing proportion of animals that are entering the water or moving around as they age or using weather-related covariates might indicate a relationship between conditions and pup availability. The detection parameters, *δ_ij_* might be similarly modeled with, for example, weather or visibility covariates or effort. Further, separate estimates can be linked through time and space in a model-based coherent framework. Currently this is often accomplished with a two-stage approach where production is estimated with the design-based method, then separate estimates are modeled through space and time using a weighted regression model (Towell et al., 2006; Johnson et al., 2013). Under the model-based framework, all of the *λ_i_* could be liked with a space-time model. Or, they could be linked using the two-stage version fitting procedure for hierarchical models proposed by Hooten et al. (2021).

## Acknowledgments and disclaimer

We would like to thank P. Conn and B. Brost for reviews of earlier versions of this paper. We would also like to thank the many field personnel in August 2016 that helped collect the fur seal mark and resight data. The findings and conclusions in the paper are those of the authors and do not necessarily represent the views of the National Marine Fisheries Service, NOAA. Reference to trade names does not imply endorsement by the National Marine Fisheries Service, NOAA.

## Author contributions

DJ, RT, and JB conceived the ideas and developed the methodology. DJ and RT formatted and analyzed the data. DJ led the writing of the manuscript. All authors contributed critically to the drafts and gave final approval for publication.

## Data availability

The fur seal data analyzed here are available at https://github.com/dsjohnson/mbpp as part of the R package containing the model fitting functions.

## References

Brintz, B., Fuentes, C., and Madsen, L. (2018). An asymptotic approximation to the n-mixture model for the estimation of disease prevalence. Biometrics, 74(4):1512–1518.

Chapman, D. G. and Johnson, A. M. (1968). Estimation of fur seal pup populations by randomized sampling. Transactions of the American Fisheries Society, 97(3):264–270.

Cochran, W. G. (1977). Sampling Techniques: 3d Ed. Wiley New York.

Cowen, L. L., Besbeas, P., Morgan, B. J., and Schwarz, C. J. (2017). Hidden Markov Models for extended batch data. Biometrics, 73(4):1321–1331.

Dail, D. and Madsen, L. (2011). Models for estimating abundance from repeated counts of an open metapopulation. Biometrics, 67(2):577–587.

Dorfman, R. (1938). A note on the delta-method for finding variance formulae. The Biometric Bulletin, 1:129–137.

Gelman, A., Lee, D., and Guo, J. (2015). Stan: A probabilistic programming language for bayesian inference and optimization. Journal of Educational and Behavioral Statistics, 40(5):530–543.

Hooten, M. B. and Hobbs, N. T. (2015). A guide to Bayesian model selection for ecologists. Ecological Monographs, 85(1):3–28.

Hooten, M. B., Johnson, D. S., and Brost, B. M. (2021). Making recursive Bayesian inference accessible. The American Statistician, 75:185–194.

Johnson, D. S., Laake, J. L., and Ver Hoef, J. M. (2010). A model-based approach for making ecological inference from distance sampling data. Biometrics, 66(1):310–318.

Johnson, D. S., Ream, R. R., Towell, R. G., Williams, M. T., and Guerrero, J. D. L. (2013). Bayesian clustering of animal abundance trends for inference and dimension reduction. Journal of Agricultural Biological and Environmental Statistics, 18(3):299–313.

Kristensen, K., Nielsen, A., Berg, C. W., Skaug, H., and Bell, B. (2016). TMB: Automatic differentiation and Laplace approximation. Journal of Statistical Software, 70(1):1–21. doi:10.18637/jss.v070.i05.

Lincoln, F. C. (1930). Calculating waterfowl abundance on the basis of banding returns. Circular 118, United States Department of Agriculture.

Link, W. A. (2013). A cautionary note on the discrete uniform prior for the binomial *N*. Ecology, 94(10):2173–2179.

Plummer, M. (2003). JAGS: A program for analysis of Bayesian graphical models using Gibbs sampling. In Proceedings of the 3rd International Workshop on Distributed Statistical Computing. Vienna, Austria.

R Core Team (2019). R: A Language and Environment for Statistical Computing. R Foundation for Statistical Computing, Vienna, Austria.

Royle, J. A. (2004). N-mixture models for estimating population size from spatially replicated counts. Biometrics, 60(1):108–115.

Schmidt, J. H., Johnson, D. S., Lindberg, M. S., and Adams, L. G. (2015). Estimating demographic parameters using a combination of known-fate and open N-mixture models. Ecology, 96(10):2583–2589.

Simpson, D. P., Rue, H., Riebler, A., Martins, T. G., and Sørbye, S. (2017). Penalising model component complexity: A principled, practical approach to constructing priors. Statistical Science, 32:1–28.

Su, Y.-S. and Yajima, M. (2015). R2jags: Using R to Run ’JAGS’. R package version 0.5-7.

Testa, J. W., editor (2018). Fur seal investigations, 2015-2016. Number NMFS-AFSC-375, 107 p. U.S. Dep. Commer., NOAA Tech. Memo. NMFS-AFSC-375, 107 p.

Towell, R., Ream, R., and York, A. (2006). Decline in northern fur seal (*Callorhinus ursinus*)pup production on the Pribilof Islands. Marine Mammal Science, 22(2):486–491.

Towell, R. G., Ream, R. R., Sterling, J. T., Bengtson, J. L., and Williams, M. (2016). Demographic studies of northern fur seals on the Pribilof Islands, Alaska, 2013-2014. In Testa, J. W., editor, Fur Seal Investigations, 2013-2014. U.S. Dept. of Commer., NOAA Tech. Memo. NMFS-AFSC-316, 124 p.

Van Dyk, D. A. and Park, T. (2008). Partially collapsed Gibbs samplers: Theory and methods. Journal of the American Statistical Association, 103(482):790–796.

York, A. and Towell, R. (1996). A new sampling design for estimating numbers of fur seal pups. In Sinclair, E. H., editor, Fur Seal Investigations, 1994. U.S. Dept. of Commer., NOAA Tech. Memo. NMFS-AFSC-69, 144 p.

